# High quality, phased genomes of *Phytophthora ramorum* clonal lineages NA1 and EU1

**DOI:** 10.1101/2021.06.23.449625

**Authors:** Nicholas C. Carleson, Caroline M. Press, Niklaus J. Grünwald

## Abstract

*Phytophthora ramorum* is the causal agent of sudden oak death in West Coast forests and currently two clonal lineages, NA1 and EU1, cause epidemics in Oregon forests. Here, we report on two high-quality genomes of individuals belonging to the NA1 and EU1 clonal lineages respectively, using PacBio long-read sequencing. The NA1 strain Pr102, originally isolated from coast live oak in California, is the current reference genome and was previously sequenced independently using either Sanger (*P. ramorum* v1) or PacBio (*P. ramorum* v2) technology. The EU1 strain PR-15-019 was obtained from tanoak in Oregon. These new genomes have a total size of 57.5 Mb, with a contig N50 length of ~3.5-3.6 Mb and encode ~15,300 predicted protein-coding genes. Genomes were assembled into 27 and 28 scaffolds with 95% BUSCO scores and are considerably improved relative to the current JGI reference genome with 2,575 or the PacBio genomes with 1,512 scaffolds. These high-quality genomes provide a valuable resource for studying the genetics, evolution, and adaptation of these two clonal lineages.

## Genome Announcement

*Phytophthora ramorum* is an invasive forest pathogen on the US West Coast causing sudden oak death (Grünwald et al. 2008; Rizzo et al. 2005). While four clonal lineages are recognized globally (Grünwald et al. 2012, 2009), only the NA1 and EU1 lineages are known to occur in US forests (Grünwald et al. 2016; Kamvar et al. 2015). The two lineages are thought to differ in aggressiveness on oaks and tanoak as well as epidemiological behavior (LeBoldus et al. 2017; Grünwald et al. 2019; O’Hanlon et al. 2017). The lineages also differ in various genomic features such as polymorphisms in presence of mitotic recombination and runs of homozygosity (ROH) (Dale et al. 2019). Novel, high quality genomes will provide a necessary resource for understanding the genetic differences among these clonal lineages.

The NA1 strain PR-102, originally isolated from coast live oak, was previously sequenced using Sanger (Tyler et al. 2006) or PacBio (Malar et al. 2019b) technology. The EU1 strain PR-15-019 was initially isolated in Oregon from tanoak bark in 2015. Strains were grown on V8 agar medium, on a sterile polycarbonate filter placed on top of the agar (Whatman Nucleopore Hydrophilic membranes, 47mm, 0.4um, GE Lifesciences) and incubated for 7 days at 20°C. High-molecular weight DNA was extracted by grinding samples in liquid N_2_ and incubating the samples in extraction buffer (20 mM EDTA, 10 mM Tris·Cl, pH 7.9, 0.5 mg/ml of both cellulase and glucanase (Sigma Aldrich Inc.,No. 219466 and G4423 respectively, St. Louis, MO), 1 % Triton^®^ X-100, 500 mM Guanidine-HCl, 200 mM NaCl) at 37°C for 2hr. Samples were then supplemented with DNase-free RNase A (20 μg/ml, Qiagen Cat. No. 19101, Germantown, MD) and incubated for 30 min at 37°C followed by proteinase-K digestion for 2h at 50°C (0.8 mg/ml, Qiagen, Cat. No. 19131). Samples were centrifuged at 12,000x *g*, clarified lysate was transferred to a Qiagen Genomic Tip 20/G kit column (Qiagen Cat. No. 10223) and DNA was separated following the manufacturer’s protocol. DNA was sequenced on a Pacific Biosciences (PacBio) Sequel I instrument at Oregon State University. The NA1 isolate PR-102 was sequenced with v2.1 chemistry on two SMRT cells, yielding 343.5x coverage. The EU1 isolate PR-15-019 was sequenced on one SMRT cell with v3.0 chemistry resulting in 184.1x coverage. All isolates were assembled using the diploid-aware SMRT tools v6.0 hierarchical genome assembly pipeline (HGAP4), which includes Arrow for polishing the assembly as a final step (Chin et al. 2013). Following HGAP, the genome was phased using Falcon-UNZIP, including a second round of polishing on the primary assembly (Chin et al. 2016). To improve the phasing, minimizing allelic variation between haplotypes falsely represented as duplicate regions, we deduplicated the primary assemblies with Purge Haplotigs (Roach et al. 2018). Briefly, the tool uses read coverage to find candidate homologous primary contigs (HPCs) and purges HPCs and places them in the alternate assembly. To determine coverage cutoffs for candidate HPCs in each genome, we mapped subreads to each assembly with minimap2 (Li 2018). Coverage histograms were evaluated independently in both genomes (*cov* options -l -h -m), since sequencing runs yielded different read abundances. Although each assembly had unique coverage distributions, the same parameters for purging (*purge* option -a 80) and clipping (*clip* options -l 30,000 -g 50,000) candidate HPCs were used on both genomes. This finalized primary assembly was scaffolded with SSPACE-LongRead (Boetzer and Pirovano 2014). Clipped sequences and purged HPCs were concatenated with alternate contigs from Falcon-Unzip and mapped against the scaffolds, using minimap2 to infer coordinates and purge_haplotigs (*place* using default options) to generate the placement files uploaded to NCBI.

BRAKER v2 was used to annotate both genomes (Lomsadze et al. 2005; Stanke et al. 2006, 2008; Ter-Hovhannisyan et al. 2008; Li et al. 2009; Barnett et al. 2011; Buchfink et al. 2015; Hoff et al. 2016, 2019; Brůna et al. 2021). Polished, deduplicated primary assemblies were soft-masked for repeats using RepeatMasker. Custom libraries for masking were generated by concatenating repeats found *de novo* by RepeatModeler in each assembly, *de novo* repeats from genomes sequenced by the Oregon State University *Phytophthora* sequencing consortium (BA Kronmiller and BM Tyler, pers. comm.), and repeats from RepBase in the *Phytophthora* lineage. As evidence for alignment, high-quality RNA-seq data for *P. ramorum* were downloaded from the NCBI SRA, trimmed with Trim Galore, and aligned to each genome using STAR (Dobin et al. 2013). Further, predicted proteomes from *P. infestans*, *P. sojae*, and *P. ramorum* were downloaded from FungiDB and aligned to each genome with Scipio (Keller et al. 2008). Hints were extracted from the Scipio alignment files using scripts from BRAKER (Keller et al. 2008). To incorporate both RNA and protein evidence, protein hints and RNA alignment files were first supplied to BRAKER using default parameters in EP mode. (Lomsadze et al. 2005; Iwata and Gotoh 2012; Gotoh et al. 2014; Buchfink et al. 2015; Brůna et al. 2020). Complementing the first run, we additionally ran BRAKER run in ETP mode to capture unique gene space. The GFFs were complemented and cleaned, including removal of redundant features, with Another Gff Analysis Toolkit (AGAT) (Dainat et al. 2020). Candidate crinklers (CRNs) and RxLR proteins were annotated from BRAKER genes complemented by six-frame translations of the genomes. CRNs were called following the HMM method in Haas et al. (2009) using confirmed CRNs from Win et al. (2006) and regular expressions in effectR (Tabima and Grünwald 2019). RxLRs were found searching for PF16180 (release 34) (Mistry et al. 2021) and the HMM method from Whisson et al. (2007) using a Python script from Cock et al. (2013).

We assembled the NA1 and EU1 genomes into 28 and 27 scaffolds, respectively, resulting in a more contiguous assembly with an N50 of >3.5 Mbp (Table 1). Fewer scaffolds capture comparable sequence space, as the total number of sequenced base pairs are larger in both new assemblies than in the first PR102 v1 assembly (Tyler et al. 2006). In addition to the improved contiguity, we report highly complete gene space with BUSCO v4 scores of 95%. Furthermore, the number of genes called are comparable to those of the Pr102 (Tyler et al. 2006) reference genome and the ND886 PacBio assembly (Table 1). Similar to the ND886 assembly, we present a phased genome, with 56.4 and 55.7 Mb of alternate sequence mapped to the NA1 and EU1 genomes respectively. These genomes provide a valuable resource for future studies on the evolution and genetics of *P. ramorum*.

**Table 1.** Genome assembly statistics of *Phytophthora ramorum* EU1 strain PR-15-019 and NA1 strain PR-102 in comparison with previous NA1 assemblies.

| Strain<br>(clonal lineage) | PR-15-019<br>(EU1) | PR-102 v3<br>(NA1) | PR-102 v2<br>(NA1) | ND886<br>(NA1) | PR-102 v1<br>(NA1) |
| --- | --- | --- | --- | --- | --- |
| Genome size (Mbp) | 58 | 58 | 70 | 60 | 67 |
| Scaffolds | 27 | 28 | 1512 | 302 | 2,576 |
| Scaffold N50 (bp) | 3,522,112 | 3,611,744 | 186,608 | 688,117 | 308,042 |
| Contigs | 52 | 71 | - | 302 | 7,589 |
| Gaps (kb) | 192.7 | 196.1 | 6.8 | 0.0 | 12,227.9 |
| Largest scaffold (Mbp) | 6.6 | 6.1 | 2.5 | 1.6 | 1.6 |
| Genes | 15,350 | 15,268 | 19,494 | 14,470 | 15,743 |
| GC Content | 0.54307 | 0.54307 | 0.54222 | 0.54451 | 0.53859 |
| Completeness (BUSCO scores)* | 93.7% (6.3%) | 94.9% (3.1%) | 93.7% (6.7%) | 88.6% (5.9%) | 93.7% (2.7%) |
| RXLR genes | 422 | 400 | 392 | 393 | 374 |
| CRN genes | 45 | 47 | 35 | 21 | 19 |
| Reference | - | - | (Malar et al. 2019b) | (Malar et al. 2019a) | (Tyler et al. 2006) |

\*BUSCO scores were calculated from the percentage of 255 Eukaryota genes of ODBv10 complete (and in duplicate).

The genome and annotation files, and raw PacBio sequencing reads, are deposited with SRA under BioProjects PRJNA738482 and PRJNA738483.

## Acknowledgements

We thank the Oregon Department of Forestry, Sarah Navarro, Jared LeBoldus, Everett Hansen and Paul Reeser for the EU1 strain, and David Rizzo for the NA1 strain. We thank Dana Gibbon, Ed Davis, and Brent Kronmiller from the Center for Genome Research and Biocomputing (CGRB) at Oregon State University for advice on PacBio sequencing, genome assembly and annotation. We thank Karan Fairchild and Valerie Fieland for technical support.

